# The role of IL-1 in adipose browning and muscle wasting in CKD-associated cachexia

**DOI:** 10.1101/2021.02.05.429984

**Authors:** Wai W Cheung, Ronghao Zheng, Sheng Hao, Zhen Wang, Alex Gonzalez, Ping Zhou, Hal M Hoffman, Robert H Mak

**Affiliations:** Pediatric Nephrology, Rady Children’s Hospital San Diego, University of California, San Diego, USA; Department of Pediatric Nephrology, Rheumatology, and Immunology, Maternal and Child Health Hospital of Hubei Province, Tongji Medical College, Huazhong University of Science and Technology, Wuhan, China; Department of Nephrology and Rheumatology, Shanghai Children’s Hospital, Shanghai Jiao Tong University, Shanghai, China; Department of Pediatrics, Shanghai General Hospital, Shanghai Jiao Tong University, Shanghai, China; Department of Pediatrics, the Second Affiliated Hospital of Harbin Medical University, Harbin, China; Department of Pediatrics, University of California, San Diego, USA

**Author notes:** These authors contributed equally to this work (Wai W Cheung, Ronghao Zheng). Correspondence: Robert H Mak, Division of Pediatric Nephrology, Rady Children’s Hospital, University of California, San Diego, 9500 Gilman Drive, MC0831, La Jolla, California 92093-0831, USA.

**Keywords:** chronic kidney disease, IL-1, cachexia, muscle wasting, adipose tissue browning

## Abstract

Cytokines such as IL-6, TNF-α and IL-1β trigger inflammatory cascades which may play a role in the pathogenesis of chronic kidney disease (CKD)-associated cachexia. CKD was induced by 5/6 nephrectomy in mice. We studied energy homeostasis in *Il1β*^−/−^/CKD, *Il6*^−/−^/CKD and *Tnfα*^−/−/^CKD mice and compared with wild type (WT)/CKD controls. Parameters of cachexia phenotype were completely normalized in *Il1β*^−/−^/CKD mice but were only partially rescued in *Il6*^−/−^/CKD and *Tnfα*^−/−^/CKD mice. We tested the effects of anakinra, an IL-1 receptor antagonist, on CKD-associated cachexia. WT/CKD mice were treated with anakinra (2.5 mg.kg.day, IP) or saline for 6 weeks and compared with WT/sham controls. Anakinra normalized food intake and weight gain, fat and lean mass content, metabolic rate and muscle function, and also attenuated molecular perturbations of energy homeostasis in adipose tissue and muscle in WT/CKD mice. Anakinra attenuated browning of white adipose tissue in WT/CKD mice. Moreover, anakinra normalized gastrocnemius weight and fiber size as well as attenuated muscle fat infiltration in WT/CKD mice. This was accompanied by correcting the increased muscle wasting signaling pathways while promoting the decreased myogenesis process in gastrocnemius of WT/CKD mice. We performed qPCR analysis for the top 20 differentially expressed muscle genes previously identified via RNAseq analysis in WT/CKD mice versus controls. Importantly, 17 differentially expressed muscle genes were attenuated in anakinra treated WT/CKD mice. In conclusion, IL-1 receptor antagonism may represent a novel targeted treatment for adipose tissue browning and muscle wasting in CKD.

## INTRODUCTION

Cachexia, characterized by muscle wasting and frailty, has been related to increased mortality rate in patients with CKD (chronic kidney disease), cancer and chronic obstructive pulmonary disease (COPD).^1,2^ A common feature of these pathological conditions is increased circulating inflammatory cytokines such as IL-1, IL-6 and TNF-α.^3–5^ Systemic inflammation is associated with exaggerated skeletal muscle wasting in CKD patients as well as animal models of CKD, suggesting anti-cytokine therapy may serve as a potential strategy to treat CKD-associated cachexia.^3^ Recent evidence in preclinical models suggests that blockade of IL-1 signaling may be a logical therapeutic target for chronic disease-associated muscle wasting. IL-1β activates NF-κβ signaling and induces expression of IL-6 and catabolic muscle atrophy atrogin-1 in C2C12 myocytes.^6,7^ Intracerebroventricular injection of IL-1β induces cachexia and muscle wasting in mice.^8^ Anakinra is an IL-1 receptor blocker for both IL-1α and IL-1β.^9^ Anakinra was FDA-approved for the treatment of rheumatoid arthritis in 2001 and is safe and an effective therapeutic option in a variety of diseases including diseases involving muscle. Duchene muscular dystrophy (DMD) is an X-linked muscle disease characterized by muscle inflammation that is associated with an increased circulating serum levels of IL-1β. Subcutaneous administration of anakinra normalized muscle function in a mouse model of DMD.^10^ Similarly, serum IL-1β is elevated in hemodialysis patients. A 4-week treatment with anakinra was shown to be safe in these patients while significantly reducing markers of systemic inflammation such as CRP and IL-6, but the effect on nutrition and muscle wasting was not established.^11^ In this study, we investigated the role of inflammatory cytokines in CKD cachexia, and evaluated the efficacy of anakinra treatment in a mouse model of CKD-associated cachexia.

## RESULTS

### Genetic deletion of IL-1β provides a better rescue of cachexia in CKD mice compared to IL-6 and TNF-α

We measured gastrocnemius muscle mRNA and protein content of IL-6, TNF-α and IL-1β in WT/CKD and compared to WT/sham mice. CKD in mice was induced by 2-stage 5/6 nephrectomy, as evidenced by higher concentration of BUN and serum creatinine (**Supplemental Table 1S**). Muscle mRNA and protein content of IL-6, TNF-α and IL-1β were significantly elevated in WT/CKD compared to WT/sham mice (**Figure 1, A-F**). In order to determine the relative functional significance of these inflammatory cytokines in CKD, we tested whether IL-6, TNF-α or IL-1β deficiency would affect the cachexia phenotype in CKD mice. WT/CKD, *Il6*^−/−^/CKD, *Tnfα*^−/−^/CKD and *Il1β*^−/−^/CKD mice were all uremic similar to WT/CKD mice (**Supplemental Table 2S**). Mice were fed *ad libitum* for 6 weeks and average daily food intake as well as final weight gain of the mice were recorded. We showed that food intake and weight gain were completely normalized in *Il1β*^−/−^/CKD mice but were only partially rescued in *Il6*^−/−^/CKD and *Tnfα*^−/−^/CKD mice (**Figure 1, G&H**). To investigate the potential metabolic effects of genetic deficiency of *Il6, Tnfα* or *Il1β* beyond its nutritional effects, we employed a pair-feeding strategy. WT/CKD mice were fed *ad libitum* and then all other groups of mice were fed the same amount of rodent diet based on the recorded WT/CKD food intake (**Figure 1I**). Despite receiving the same amount of total calorie intake as other groups of mice, cardinal features of cachexia phenotype such as decreased weight gain, decreased fat mass content and hypermetabolism (manifested as elevated oxygen consumption), decreased lean mass content and reduced muscle function (grip strength) were evident in WT/CKD mice (**Figure 1, J to N**). Importantly, the cachexia phenotype was completely normalized in *Il1β*^−/−^/CKD mice relative to *Il1β*^−/−^/sham mice whereas *Il6*^−/−^/CKD and *Tnfα*^−/−^/CKD mice still exhibited some degree of cachexia.

**Figure 1:**
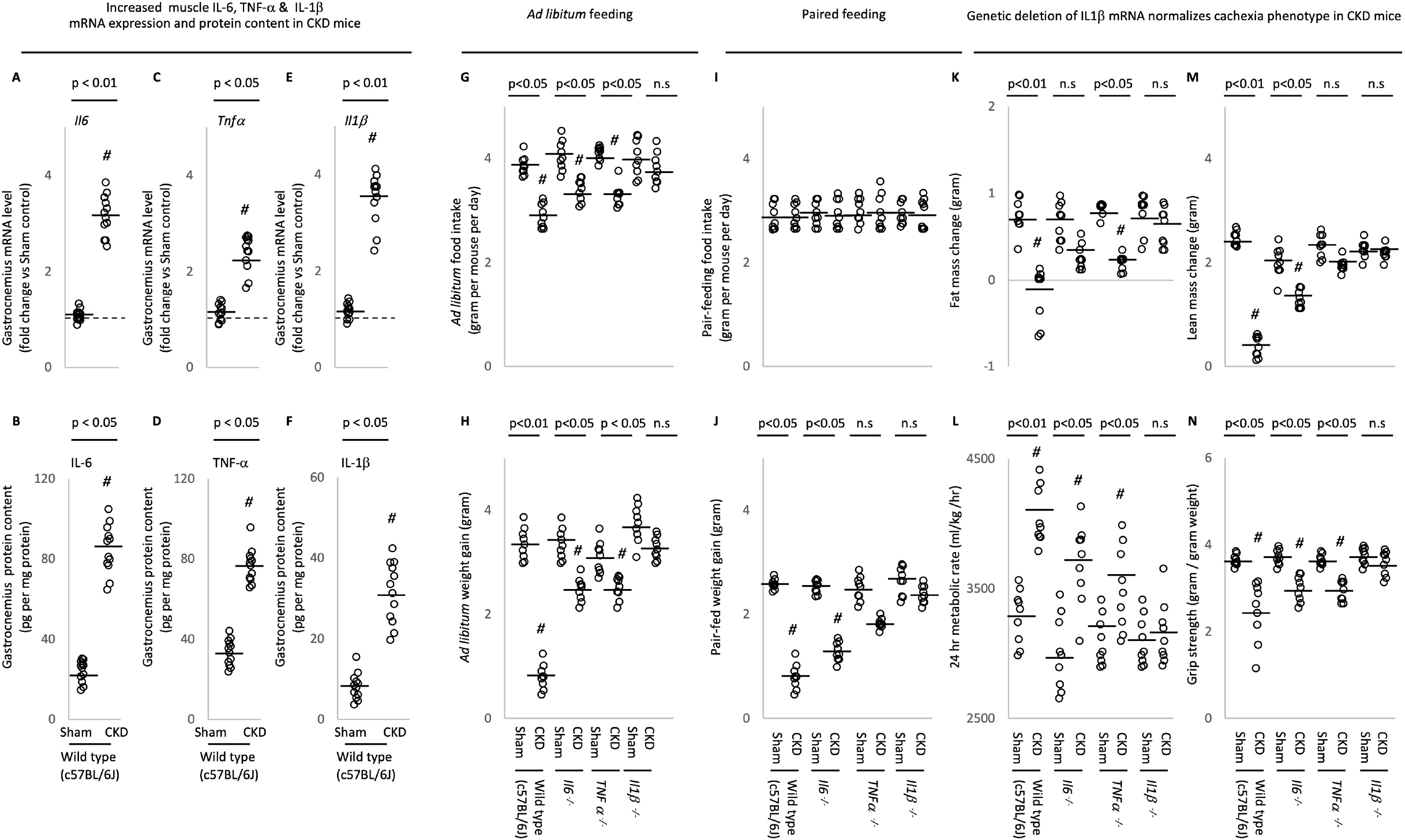
Increased muscle mRNA and protein content of IL-6, TNF-α and IL-1β in CKD mice and genetic depletion of IL-1β provides a better rescue of cachexia in CKD mice compared to IL-6 and TNF-α deficiency. Results of 3 experiments were shown. For the 1^st^ experiment, CKD was induced by 5/6 nephrectomy in WT mice and sham operation was performed in WT control mice. Gene expression and protein content of pro-inflammatory cytokine IL-6, TNF-α and IL-1β in gastrocnemius muscle in WT/CKD and WT/sham mice was performed. Data are expressed as mean ± SEM and results of WT/CKD mice were compared to WT/Sham mice (A to F). For the 2^nd^ experiment, 4 groups of mice were included: WT/CKD and WT/sham, *Il6*^−/−^/CKD and *Il6*^−/−^/sham, *Tnfα*^−/−^/CKD and *Tnfα*^−/−^/sham, *Il1β*^−/−^/CKD and *Il1β*^−/−^/sham. All mice were fed *ad libitum* and food intake as well as weight gain in mice were recorded (G & H). For the 3^rd^ experiment, we investigated the metabolic benefits of genetic depletion of IL-6, TNF-α and IL-1β in CKD mice beyond the nutritional stimulation by employed a pair-feeding strategy. Four groups of mice are WT/CKD and WT/sham, *Il6*^−/−^/CKD and *Il6*^−/−^/sham, *Tnfα*^−/−^/CKD and *Tnfα*^−/−^/sham, *Il1β*^−/−^/CKD and *Il1β*^−/−^/sham. WT/CKD mice were fed *libitum* whereas other groups of mice were pair-fed to that of WT/CKD mice (I). Weight gain, fat and lean content, 24-hr oxygen consumption and *in vivo* muscle function (grip strength) was measured in mice (J to N). For the 2^nd^ and 3^rd^ experiments, data are expressed as mean ± SEM and results of WT/CKD, *Il6*^−/−^/CKD, *Tnfα*^−/−^/CKD and *Il1β*^−/−^/CKD mice were compared to WT/sham, *Il6*^−/−^/sham, *Tnfα*^−/−^/sham and *Il1β*^−/−^/sham, respectively.

### Anakinra attenuates cachexia in CKD mice

In order to test the efficacy of anakinra in murine CKD, WT/CKD and WT/sham mice were treated with anakinra or vehicle for 6 weeks. All mice were fed *ad libitum*. Food intake and weight gain were normalized in anakinra treated WT/CKD mice (**Figure 2, A&B**). We also investigated the beneficial effects of anakinra in WT/CKD mice beyond appetite stimulation and its consequent body weight gain through the utilization of a pair-feeding approach. Daily *ad libitum* food intake for WT/CKD mice treated with vehicle was recorded. Following that, anakinra treated WT/CKD mice were food restricted such that the mice ate an equivalent amount of food as vehicle treated WT/CKD mice (**Figure 2C**). Serum and blood chemistry of mice were measured (**Supplemental Table 3S**). Anakinra normalized weight gain, metabolic rate, fat and lean mass content, gastrocnemius weight, as well as *in vivo* muscle function (rotarod and grip strength) in WT/CKD mice (**Figure 2, F to J**).

**Figure 2:**
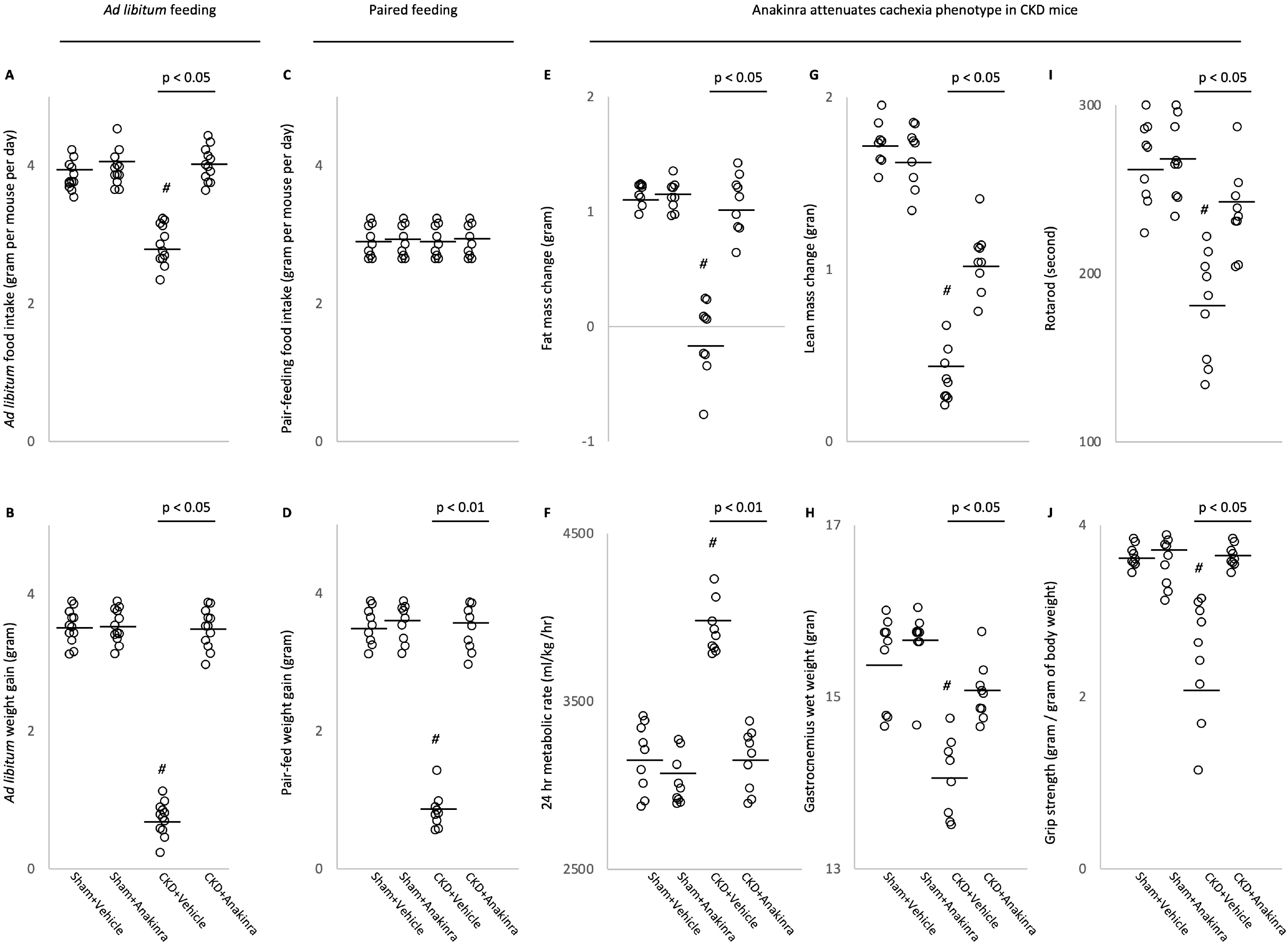
Anakinra attenuates cachexia in CKD mice. WT/CKD and WT/sham mice were treated with anakinra (2.5 mg.kg.day, IP, daily) or normal saline as a vehicle control for 6 weeks. Mice were fed *ad libitum* and food intake and weight gain in mice were recorded (A & B). To assess the beneficial effects of anakinra beyond its nutritional effects, we employed a pair-feeding strategy. WT/CKD mice treated with vehicle were given an *ad libitum* amount of food whereas other groups of mice were given an equivalent amount of food (C). Weight change, fat and lean mass content, 24-hr oxygen consumption and *in vivo* muscle function (grip strength) was measured in mice (D to J). Data are expressed as mean ± SEM. Results of WT/CKD+Vehicle were compared to WT/sham+Vehicle and WT/CKD+Anakinra were compared to WT/sham+Anakinra, respectively. In addition, results of WT/CKD+Anakinra were also compared to WT/CKD+Vehicle. *#* p < 0.05.

### Anakinra normalizes energy homeostasis in skeletal and adipose tissue in CKD mice

Adipose tissue (inguinal WAT and intercapsular BAT) and muscle protein content of UCPs was significantly increased in WT/CKD mice compared to control mice (**Figure 3, A, C & E**). In contrast, ATP content in adipose tissue and skeletal muscle was significantly decreased in WT/CKD mice compared to control mice (**Figure 3, B, D & F**). Anakinra normalized UCPs and ATP content of adipose tissue and muscle in WT/CKD mice.

**Figure 3:**
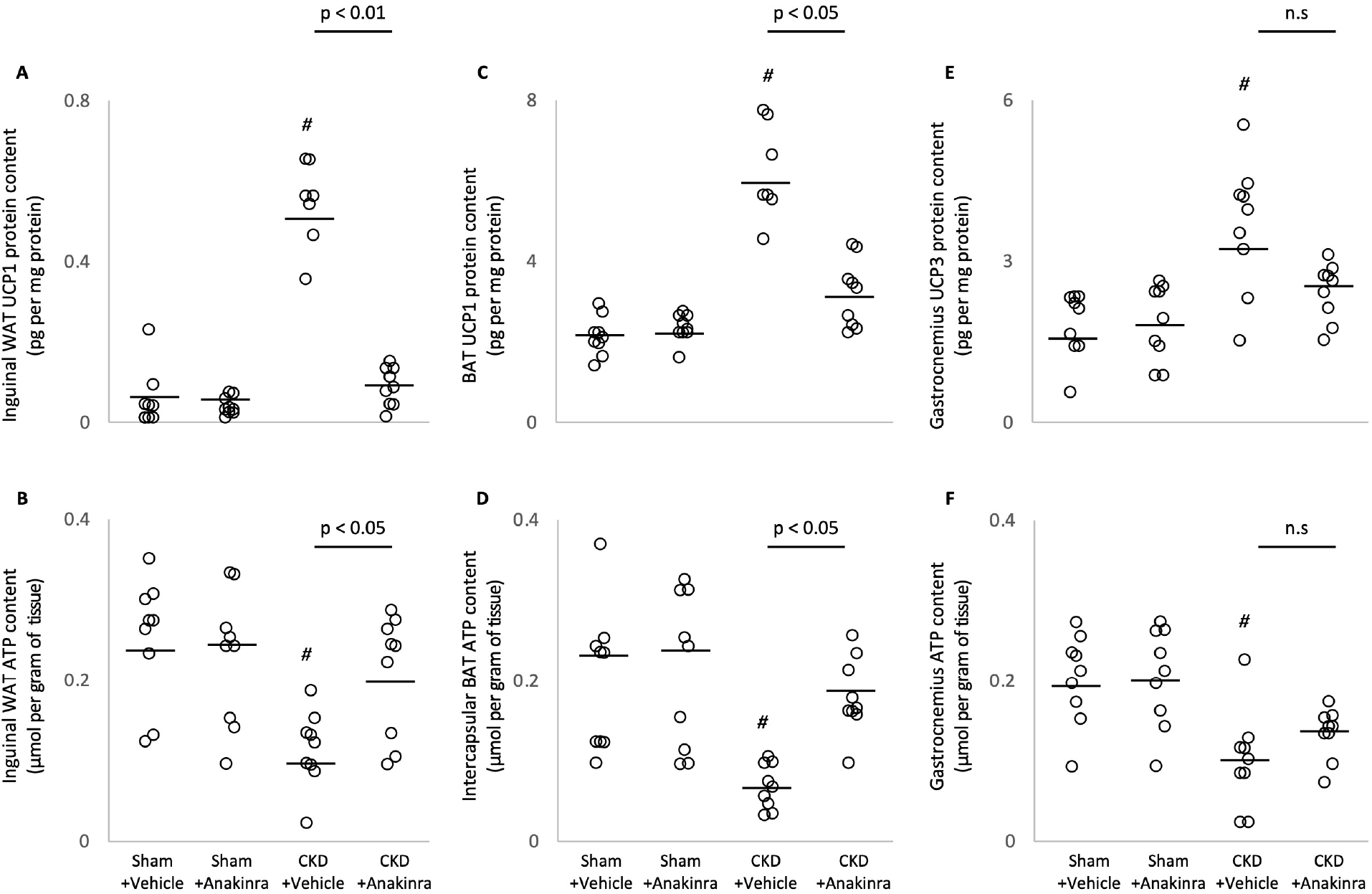
Anakinra ameliorates energy homeostasis in skeletal and adipose tissue in CKD mice. UCPs and ATP protein content in adipose tissue (inguinal white adipose tissue and brown adipose tissue) and gastrocnemius muscle were measured. Final results were expressed in arbitrary units, with one unit being the mean level in WT/sham+Vehicle mice. Results are analyzed and expressed as in Figure 2.

### Anakinra ameliorates browning of white adipose tissue in CKD mice

Expression of mRNA of beige adipose cell surface markers (*CD137, Tmem26, and Tbx1*) was elevated in inguinal WAT in WT/CKD mice relative to sham control mice (**Supplemental Figure 1**). Moreover, inguinal WAT protein content of CD137, Tmem26 and Tbx1 were significantly increased in WT/CKD mice compared to control mice (**Figure 4, A to C**). This was accompanied by an increase UCP-1 protein content in inguinal WAT shown in **Figure 3A**, which is a biomarker for beige adipocyte, and usually undetectable in WAT. Anakinra normalized protein content of UCP-1 and attenuated protein content of CD137, Tmem26 and Tbx1 in WT/CKD mice relative to vehicle treated control mice (**Figure 3A and Figure 4, A to C**). Furthermore, anakinra ameliorated expression of important molecules involved in browning of adipose tissue in WT/CKD mice. Activation of Cox2/Pgf2α signaling pathway and chronic inflammation stimulate biogenesis of WAT browning. Significantly increased inguinal WAT mRNA expression (**Supplemental Figure 1**) and protein content of Cox2 and Pgf2α was found in WT/CKD mice compared to controls (**Figure 4, D and E**). Moreover, inguinal WAT levels of toll-like receptor 2 (Tlr2) as well as myeloid differentiation primary response 88 (MyD88) and TNF associated factor 6 (Traf6) were upregulated in WT/CKD mice compared to control mice (**Supplemental Figure 1 and Figure 4, F to H**). Anakinra partially or fully normalized inguinal WAT expression of Cox2/Pgf2α as well as the toll-like receptor pathway in WT/CKD mice relative to vehicle treated controls.

**Figure 4:**
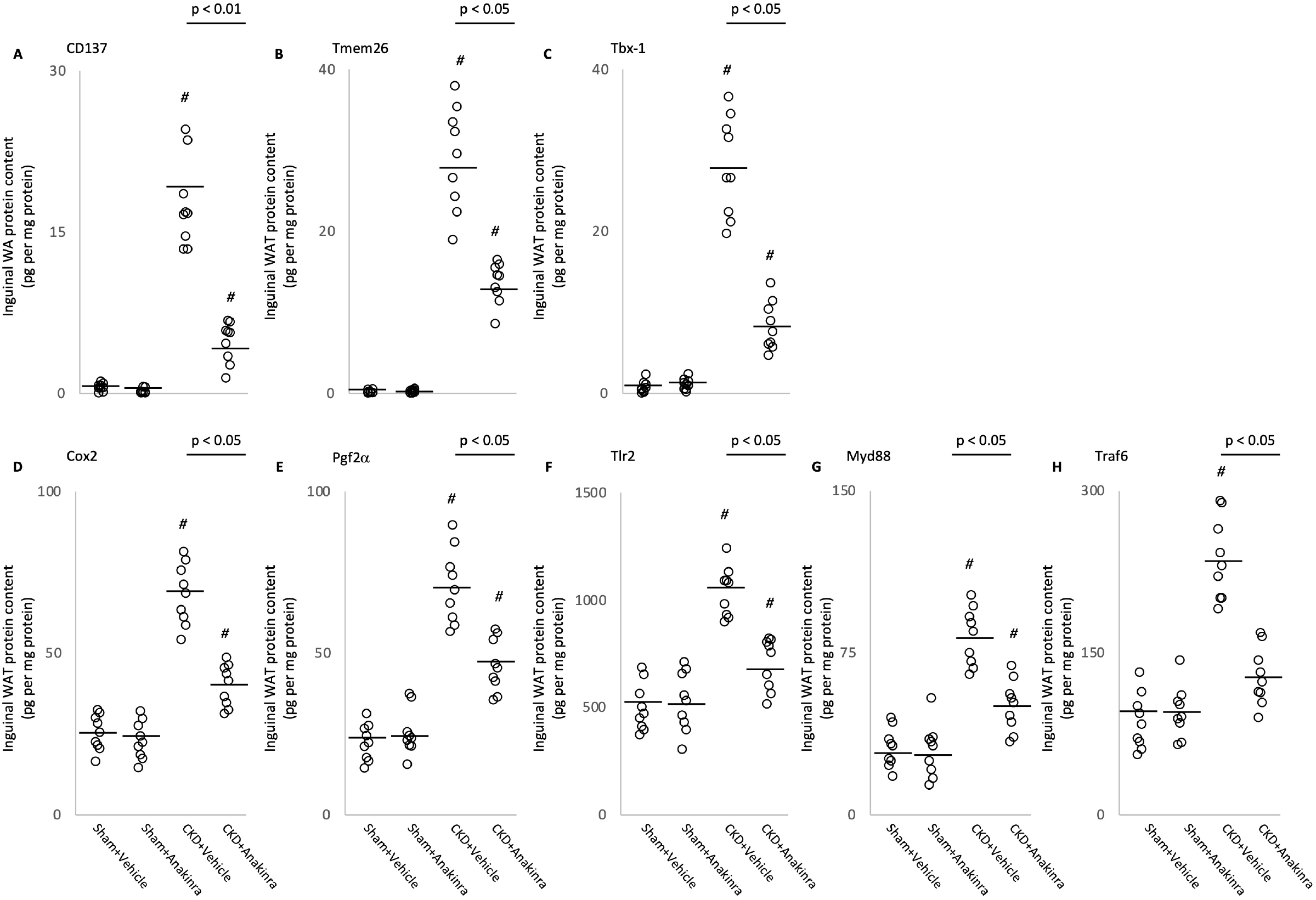
Anakinra attenuates adipose tissue browning in CKD mice. Protein content of beige adipocyte markers (CD137, Tmem26 and Tbx-1) in inguinal white adipose tissue was measured. In addition, protein content of Cox2 signaling pathway (Cox2 and Pgf2a) and toll like receptor pathway (Tlr2, MyD88 and Traf6) in inguinal white adipose tissue was measured. Results are analyzed and expressed as in Figure 2.

### Anakinra attenuates signaling pathways implicated in muscle wasting in CKD mice

The ratio of muscle phospho-NF-κB p50 (phosphorylated Ser337), phospho-NF-κB p65 (phosphorylated Ser536) relative to total NF-κB p50 and p65 content, respectively, was significantly increased in WT/CKD relative to sham control mice (**Figure 5, A & B**). Anakinra did not change protein content of muscle phospho-Iκκα relative to total Iκκα in WT/CKD mice (**Figure 5C**). Muscle phospho-AKT (pS473), phospho-ERK 1/2 (Thr202/Tyr204), phospho-JNK (Thr183/Tyr185) and phosphor-p38 MAPK content (Thr180/Tyr182) was elevated in WT/CKD mice compared to sham control mice (**Figure 5, D to G**). Importantly, anakinra attenuated or normalized phosphorylation of muscle phospho-NF-κB p50 and p65 content as well as phospho-AKT, phospho-ERK 1/2, phospho-JNK, phospho-p38 MAPK in WT/CKD mice. Administration of anakinra improved muscle regeneration and myogenesis by decreasing mRNA expression of negative regulators of skeletal muscle mass (Atrogin-1, Murf-1 and Myostatin) (**Figure 5, H to J**) as well as increasing muscle mRNA expression of pro-myogenic factors (IGF-1, Pax-7, MyoD and Myogenin) in WT/CKD mice (**Figure 5, K to N**).

**Figure 5:**
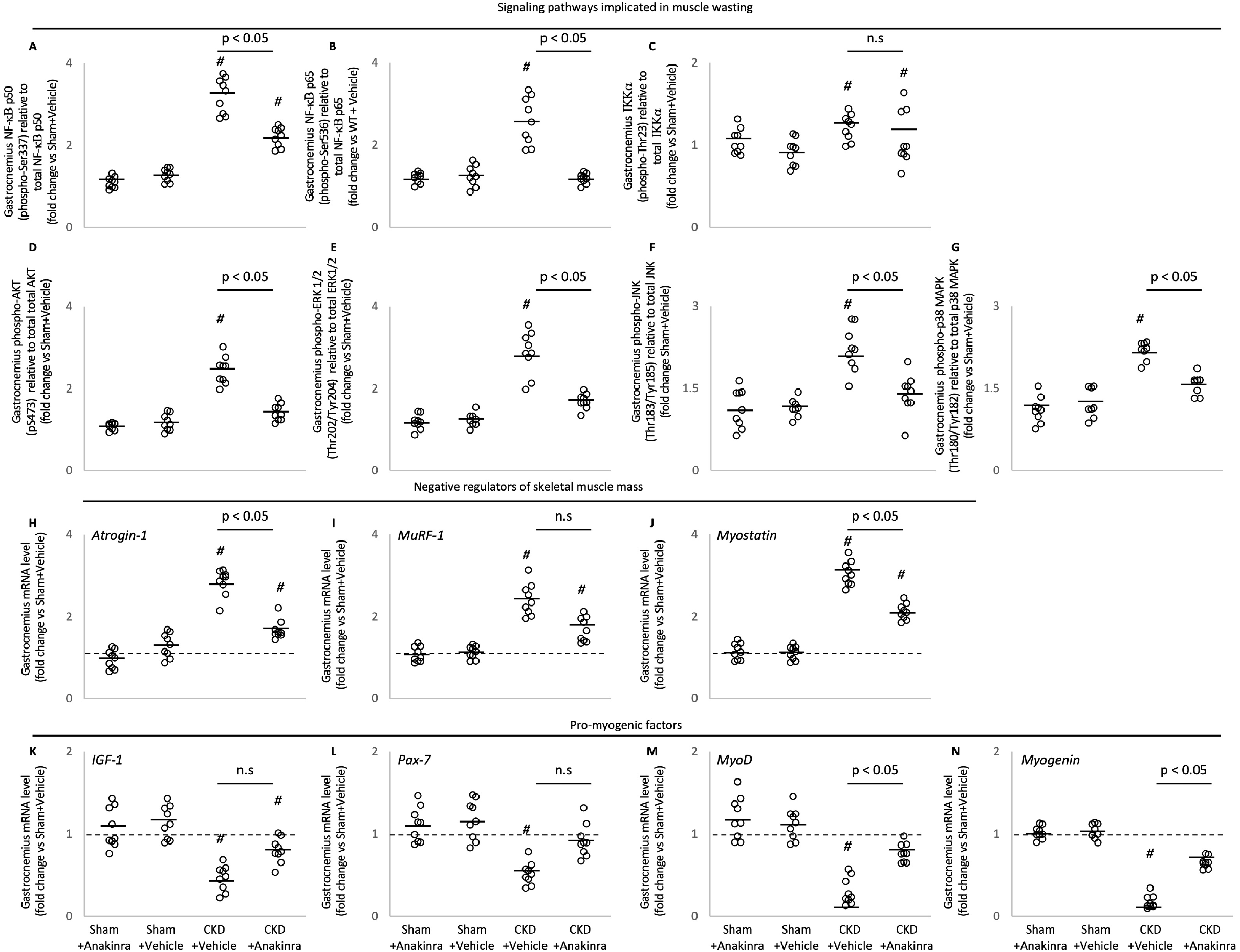
Anakinra attenuates signaling pathways implicated in muscle wasting in CKD mice. Gastrocnemius muscle relative phosphorylated NF-κB p50 (Ser337) / total p50 ratio, NF-κB p65 (Ser536) / total p65 ratio and Iκκ*a* (Thr23) / total Iκκ*a* ratio as well as muscle relative phospho-Akt (pS473) / total Akt ratio, ERK 1/2 (Thr202/Tyr204) / total ERK 1/2 ratio, JNK (Thr183/Tyr185) / total JNK ratio, p38 MAPK (Thr180/Tyr182) / total p38 MAPK ratio in mice. In addition, gastrocnemius muscle expression of interested genes in mice was measured by qPCR. Final results were expressed in arbitrary units, with one unit being the mean level in WT/sham+Vehicle mice. Results are analyzed and expressed as in Figure 2.

### Anakinra improves muscle fiber size and attenuates muscle fat infiltration in CKD mice

We evaluated the effect of anakinra on skeletal muscle morphology in WT/CKD mice. Representative results of muscle sections are illustrated (**Figure 6A**). Anakinra attenuated average cross-sectional area of gastrocnemius in WT/CKD mice (**Figure 6B**). We also evaluated fat infiltration of the gastrocnemius muscle in WT/CKD mice. Representative results of Oil Red O staining of muscle section are shown (**Figure 6C)**. We quantified intensity of muscle fat infiltration in mice and observed that anakinra attenuated muscle fat infiltration in WT/CKD mice (**Figure 6D**).

**Figure 6:**
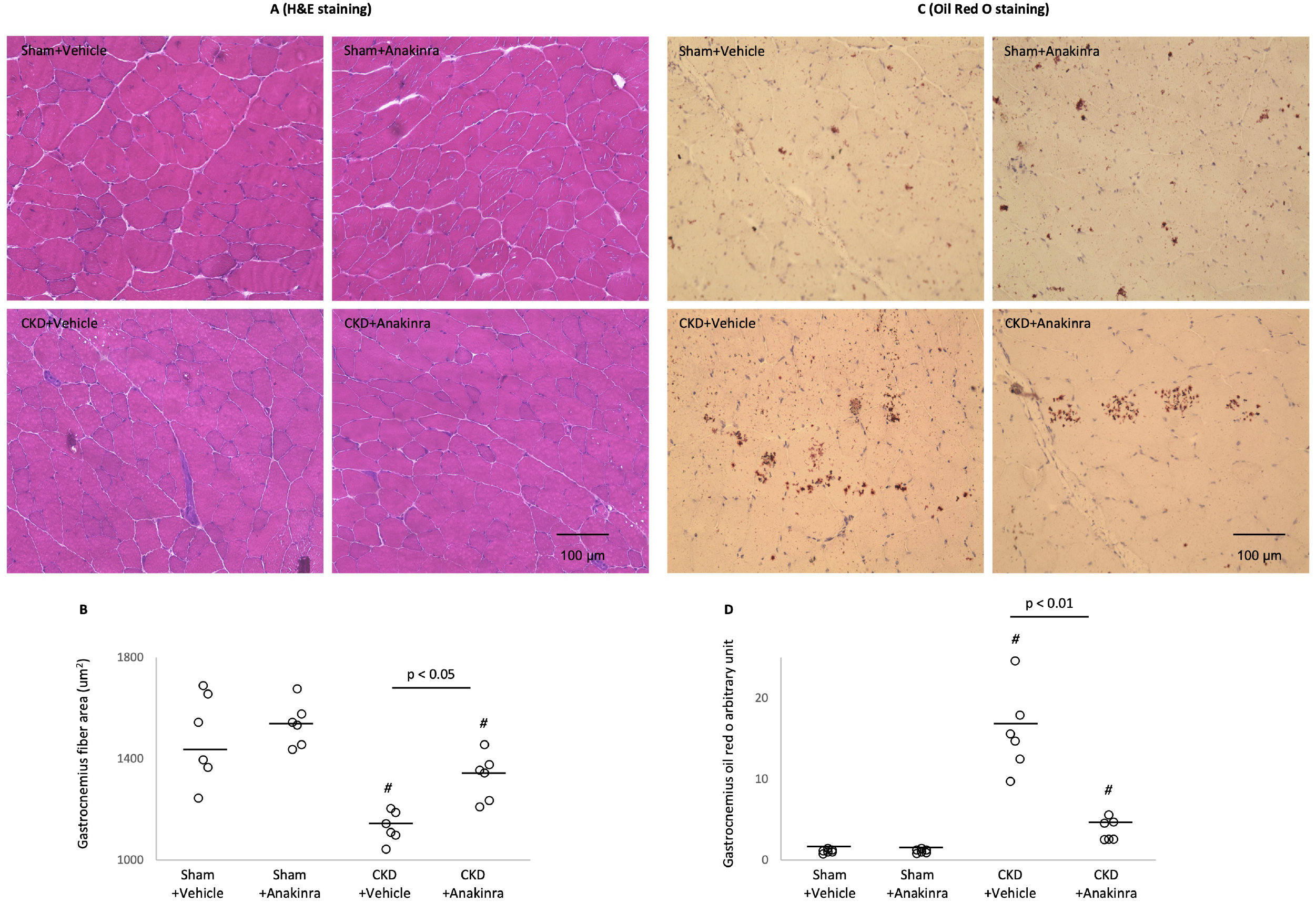
Anakinra normalizes muscle fiber size and attenuates muscle fat infiltration in CKD mice. Representative photomicrographs of gastrocnemius with H&E staining (A). Average gastrocnemius cross-sectional area was measured (B). Visualization of quantification of fatty infiltration by Oil Red O analysis in gastrocnemius muscle (C and D). Final results were expressed in arbitrary units, with one unit being the mean staining intensity in vehicle-treated WT/sham mice. Difference among various groups of mice were analyzed as in Figure 2.

### Molecular mechanism by RNAseq analysis

We studied differential expression of gastrocnemius mRNA between 12-months old WT/CKD mice and WT control mice using RNAseq analysis.^12^ Ingenuity Pathway Analysis enrichment tests have identified top 20 differentiated expressed muscle genes in WT/CKD mice versus WT control mice. We performed qPCR analysis for top 20 differentially expressed muscle genes in the present study. Muscle gene expression of Sell, Tnni1 and Tnnt1 remained significantly elevated in WT/CKD mice (**Supplemental Figure 2**). Importantly, anakinra normalized (*Atf3, Atp2a2, Fhl1, Fosl2, Gng2, Itpr1, Lamc3, Mafb, Maff, Myl2, Nlrc3, Pthr1, Tnnc1, Tpm3* and *Ucp2*) and attenuated (*Csrp3* and *Cyfip2*) muscle gene expression in WT/CKD mice relative to WT sham mice (**Figure 7**). We summarize the functional significance of these 17 differentially expressed muscle genes that were normalized or attenuated in anakinra treated WT/CKD mice. (**Table 1**).

**Figure 7:**
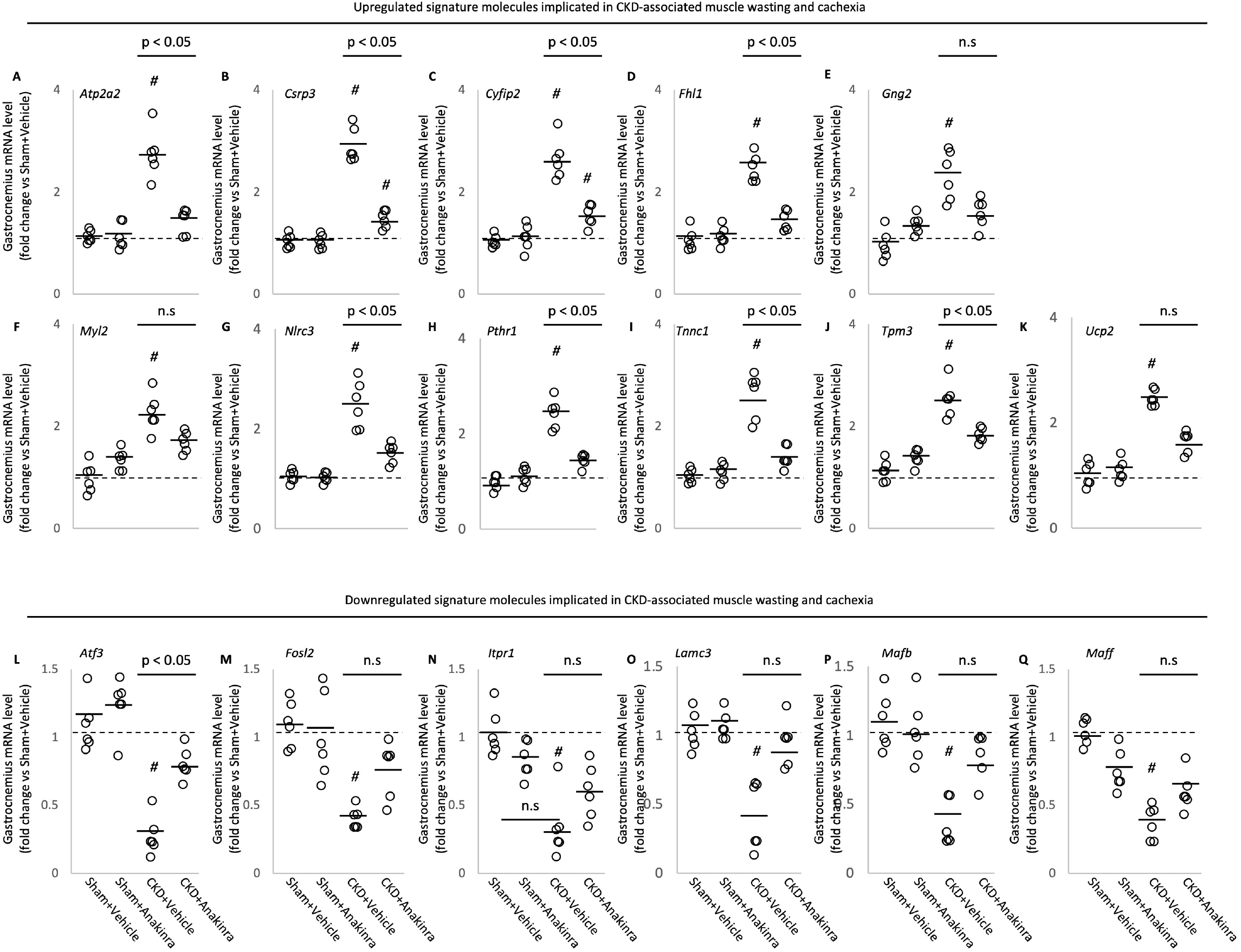
Anakinra attenuates expression of top 17 differentiated expression gastrocnemius muscle genes in CKD mice. Gastrocnemius muscle expression of interested genes in mice was measured by qPCR. Final results were expressed in arbitrary units, with one unit being the mean level in vehicle-treated WT/sham mice. Results are analyzed and expressed as in Figure 2.

**Table 1:** Anakinra normalizes or attenuates expression of important muscle genes that have been implicated in muscle wasting in CKD mice. Previously, we studied differential expression of gastrocnemius mRNA between 12-months old WT/CKD mice and WT sham mice using RNAseq analysis.^12^We focus on pathways related to energy metabolism, skeletal and muscular system development and function, nervous system development and function as well as organismal injury and abnormalities. We performed qPCR analysis for top 20 differentially expressed muscle genes in the present study. Importantly, anakinra normalized (*Atf3, Atp2a2, Fhl1, Fosl2, Gng2, Itpr1, Lamc3, Mafb, Maff, Myl2, Nlrc3, Pthr1, Tnnc1, Tpm3, Ucp2*) and attenuated (*Csrp3, Cyfip2*) muscle gene expression in WT/CKD mice relative to WT sham mice. Functional significance of each of these 17 differentially expressed muscle genes is listed.

| Upregulated differential expressed genes | Functional significance & references |
| --- | --- |
| *Atp2a2* | associated with myogenic differentiation in slow-twitch muscle fiber <sup>56</sup> |
| *Csrp3* | associated with skeletal muscle dystrophy <sup>46</sup> |
| *Cyfp2* | associated with muscle wasting <sup>47</sup> |
| *Fhl1* | activates myostatin signaling and promotes atrophy in skeletal muscle <sup>48</sup> |
| *Gng2* | biomarker of fatty infiltration in muscle <sup>60</sup> |
|  | associated with adipocyte morphology and metabolic derangements <sup>60</sup> |
| *Myl2* | associated with muscle wasting by inhibiting myoblast proliferation <sup>50</sup> |
| *Nlrc3* | implicated in skeletal muscle wasting by inhibiting cell proliferation and promoting cell apoptosis <sup>51</sup> |
| *Pthr1* | associated with muscle fatigue, cardiovascular pathology and hyperparathyroidism <sup>55</sup> |
|  | implicated in parathyroid-hormone-dependent skeletal muscle mass metabolism <sup>52</sup> |
| *Tnnc1* | regulates straited muscle contraction <sup>53</sup> |
|  | implicated in cardiomyopathy pathogenesis and age-related skeletal muscle wasting <sup>53</sup> |
| *Tpm3* | promotes slow myofiber hypotrophy and associated with generalized muscle weakness <sup>54</sup> |
| *Ucp2* | regulates body metabolism and implicated in skeletal muscle wasting <sup>61,62</sup> |
| Downregulated differential expressed genes | Functional significance & references |
| *Atf3* | impairs motor neuron survival and muscle innervation <sup>57</sup> |
| *Fosl2* | implicated in FoxO-dependent gene network in muscle wasting <sup>53</sup> |
| *Itpr1* | impairs muscle regeneration <sup>58</sup> |
| *Lamc3* | impairs muscle regeneration and induces muscle dystrophy <sup>59</sup> |
| *Mafb* | implicated with Duane retraction syndrome in patients with focal segmental glomerulosclerosis <sup>64</sup> |
| *Maff* | implicated in Prdm4-induced white adipose tissue browning <sup>63</sup> |

## DISCUSSION

Inflammation plays a major role in cachexia from different underlying diseases. Specifically, the inflammatory cytokines, IL-6, TNF-α and IL-1 have been implicated in CKD-associated cachexia. Our studies using specific cytokine deficient mice and the IL-1 targeted therapy anakinra suggest that IL-1 is the most important cytokine in CKD associated cachexia. We showed that anakinra normalized food intake and weight gain, fat and lean mass content, metabolic rate and muscle function in CKD mice. Moreover, anakinra attenuated browning of white adipose tissue in CKD mice. Furthermore, anakinra normalized gastrocnemius weight and fiber size as well as attenuated muscle fat infiltration in CKD mice. This was accompanied by correcting the increased muscle wasting signaling pathways while promoting the decreased myogenesis process in gastrocnemius of CKD mice. Together, our results suggest that anakinra may be an effective targeted treatment approach for cachexia in patients with CKD.

Peripheral or central administration of IL-1β suppresses food intake, activates energy metabolism and reduces weight gain in experimental animals.^8,13^ IL-1β signals through the appetite-regulating neuropeptides, such as leptin resulting in appetite suppresion.^14^ Previously, we have demonstrated that elevated circulating level of leptin through the activation of melanocortin receptor 4 induces CKD-associated cachexia.^15^ In this study, we showed that skeletal muscle IL-1β level was significantly increased in CKD mice, and *Il1β*^−/−^/ mice had attenuated CKD induced cachexia (**Figure 1**). Furthermore, we showed that anakinra improved anorexia and normalized weight gain in WT mice with CKD (**Figure 2, A and B**). Our results further highlight the beneficial effects of anakinra beyond food stimulation and accompanied weight gain. In pair-fed studies in which CKD and control mice were fed the same amount of food, cachexia was attenuated in CKD mice treated with anakinra than control mice (**Figure 2D**).

The beneficial effects of administration of anakinra in CKD mice were in agreement with human data in the context of CKD-associated cachexia. Deger *et al* have shown that systemic inflammation, as assessed by increased serum concentration of CRP, is a strong and independent risk factor for skeletal muscle wasting in CKD patients.^3^ Their results provide rationale for further studies using anti-cytokine therapies for patients with CKD. Administration of anakinra reduced inflammatory response in CKD patients as reflected by significant decreases in plasma concentration of inflammatory biomarkers CRP and IL-6.^11^ Subsequent study by the same group also showed that blockade of IL-1 significantly reduced inflammatory status (decreased plasma concentration of IL-6, TNF-α and Nod-like receptor protein 3) as well as improved antioxidative property (increased plasma concentration of superoxide) in patients with stages 3-5 CKD.^16^ However the impact of this anti-IL-1 therapy on wasting and cachexia in CKD patients has not been established.

Loss of adipose tissue is a crucial feature of cachexia and is associated with increased lipolysis or decreased adipogenesis. Adipogenesis, the formation of adipocytes from stem cells, is important for energy homeostasis and is involved in processing triglycerol, the largest energy reserve in the body.^17^ IL-1β inhibits adipogenesis as suggested by the finding that potential of adipogenic progenitor cells isolated from patients with DMD are significantly reduced when co-cultured with IL-1β-secreting macrophages.^18^ We showed that anakinra normalized fat content in CKD mice (**Figure 2E**). The basal metabolic rate accounts for up to 80% of the daily calorie expenditure by individual.^19^ Skeletal muscle metabolism is a major determinant of resting energy expenditure.^20,21^ IL-1β increases basal metabolic rate (as represented by an increase in resting oxygen consumption) in a dose-dependent manner.^22^ In our study, anakinra normalized the increased 24-hr metabolic rate in CKD mice (**Figure 2F**).

Adipose tissue UCP1 expression is essential for adaptive adrenergic non-shivering thermogenesis and muscle UCP3 level controls body metabolism.^23^ The energy generated when dissipating the proton gradient via upregulation of UCPs is not used for cellular ATP production or other biochemical processes but instead to produce heat.^24,25^ We showed that anakinra normalized muscle and adipose tissue ATP and UCPs content in CKD mice (**Figure 3**). Blockade of IL-1 receptor signaling may also mitigate the metabolic dysfunction through leptin signaling. Infusion of leptin increased UCPs expression in skeletal muscle and adipose tissue.^26,27^

Adipose tissue browning is associated with profound energy expenditure and weight loss in patients with CKD-associated cachexia.^28–30^ We previously demonstrated adipose tissue browning in CKD mice (as evidenced by the detection of inguinal WAT UCP1 protein and increased expression of beige adipose cell markers CD137, Tmem26 and Tbx1).^12,31^ Activation of the Cox2 signaling pathway and chronic inflammation induce adipose tissue browning. Cox2 is a downstream effector of β-adrenergic signaling and induces biogenesis of beige cells in WAT depots.^32^ In this study, we showed that anakinra attenuated inguinal WAT protein and mRNA content of Cox2 and Pgf2a level in CKD mice as well as normalized key inflammatory molecules (Tlr2, MyD88 and Traf6) involved in adipose tissue browning in CKD mice (**Figure 4 and Supplemental Figure 1**). Recent data suggest that IL 1β signaling mediates adipocyte browning and thermogenesis via regulation of mitochondrial oxidative responses in both cultured human and animal adipocytes.^33^

We investigated the effects of anakinra on muscle wasting in CKD mice. Lean mass content and gastrocnemius muscle weight was significantly decreased in CKD (**Figure 2, G and H**). We showed that anakinra normalized lean mass content, gastrocnemius weight as well as muscle function in CKD mice. We then investigated the impact of anakinra on muscle fiber morphology in CKD mice. Anakinra normalized average cross-sectional area of gastrocnemius muscle in CKD mice (**Figure 6, A and B**). Muscle fat infiltration is a significant predictor of both muscle function and mobility function across a wide variety of comorbid conditions such as diabetes, spinal cord injury and kidney disease.^34^ Intramuscular fat infiltration has been implicated in metabolic dysfunction such as insulin resistance. Muscle adipose tissue may release pro-inflammatory cytokines within the muscle and impair the local muscle environment, impair blood flow or increase the rate of lipolysis within skeletal muscle resulting in an increased concentration of glucose within the skeletal muscle itself followed by insulin resistance.^34^ Patients with CKD exhibit low muscle insulin sensitivity even at the onset of renal dysfunction. This uremic muscle resistance has been linked to disturbed protein metabolism and to the loss of skeletal muscle function and mass.^35^ We showed that anakinra attenuated muscle fat infiltration in CKD mice (**Figure 6, C and D**).

IL-1 has been shown to stimulate the expression of catabolic genes.^4,9,36^ Several important signaling pathways regulate skeletal muscle mass metabolism. Upregulation of Akt/mTOR pathway stimulates skeletal muscle hypertrophy and atrophy.^37^ Inhibition of ERK signaling attenuated muscle wasting via promoting myogenesis in tumor bearing mice.^38,39^ JNK signaling is activated in mouse model of pancreatic cancer cachexia and inhibition of JNK signaling improves body weight and muscle strength (grip strength) in tumor-bearing mice.^40^ MAPK are a family of protein phosphorylating enzymes that regulate a diverse aspect of cellular responses including skeletal muscle regeneration and differentiation.^41–43^ Emerging evidence suggests that NF-κB is one of most important signaling pathways linked to the loss of skeletal muscle mass in various physiological and pathophysiological conditions. Activation of NF-κB in skeletal muscle leads to degradation of muscle proteins, induces inflammation and fibrosis, and blocks the regeneration of myofibers.^44,45^ In this study, we showed that anakinra normalized or reduced phosphorylation of muscle Akt, ERK, JNK, MAPK, NF-κB p50 and p65 content in CKD mice (**Figure 5, A&B, D to G**). This was accompanied by decreasing the gene expression of negative regulators of skeletal muscle mass (*Atrogin-1, Murf-1* and *Myostatin*) while increasing the gene expression of pro-myogenic factors (*IGF-1, Pax-7, MyoD* and *Myogenin*) in CKD mice (**Figure 5, H to N**).

Recently, we identified a gene expression signature by RNA sequence analysis in muscle of CKD mice compared to control mice.^12^ We performed qPCR analysis for the top 20 differentially expressed muscle genes in the present study. Anakinra treatment normalized (*Atf3, Atp2a2, Fhl1, Fosl2, Gng2, Itp1, Lamc3, Mafb, Maff, Myl2, Nlcr3, Pthr1, Tnnc1, Tpm3, Ucp2*) and attenuated (*Csrp3, Cyfip2*) gene expression in muscle from CKD mice (**Figure 7**). Aberrant gene expression of *Csrp3, Cyfip2, Fhl1, Fosl2, Myl2, Nlcr3, Pthr1, Tnnc1* and *Tpm3* have been implicated in muscle atrophy and muscle weakness.^46–54^ Parathyroid hormone (PTH) and its receptor may mediate the crosstalk between adipose tissue and muscle in CKD cachexia. Parathyroid hormone 1 receptor (PTH1R) functions as a receptor for Parathyroid hormone (PTH) and PTH-related peptide (PTHrP).^55^ PTH and PTHrP, which signal through the same receptor PTH1R, induce adipose tissue and muscle wasting in murine models of cancer and CKD.^28,30^ Increased expression of Pthr1 has been associated with muscle fatigue, cardiovascular pathology and hyperparathyroidism as well as skeletal muscle wasting.^52,55^ CKD mice in this study had elevated circulating PTH levels but anakinra did not normalize serum PTH levels in CKD mice (**Supplemental Table 3S**), suggesting that PTH/PTHrP pathway may not be the only mediator of crosstalk between adipose tissue and muscle in CKD-associated cachexia. We did find that muscle Pthr1 gene expression was significantly increased in CKD mice and anakinra normalized upregulated muscle Pthr1 expression in CKD mice (**Figure 7H**). Increased expression of *Atp2a2* and decreased expression of *Atf3, Itpr1* and *Lamc3* have been associated with impaired motor neuron survival and muscle innervation, reduced myogenic differentiation and regeneration.^55–59^ Moreover, increased *Gng2* expression is a biomarker of fatty infiltration in muscle and increased muscle Gng2 expression has been associated with aberrant adipocyte morphology and metabolic derangements in various metabolic diseases.^60^ Increased Ucp2 expression stimulates body metabolism and promotes skeletal muscle wasting.^61,62^ Results also suggest that decreased expression of *Maff* is implicated in white adipose tissue browning.^63^ Interestingly, aberrant expression of *Mafb* has been associated with Duane retraction syndrome in patient with focal segmental glomerulosclerosis.^64^ In **Table 1**, we list the functional significance of each of the 17 differentially expressed muscle genes that has been normalized or attenuated in anakinra treated CKD mice relative to control mice.

In summary, we report that IL-1 antagonism and specific pharmacological blockade using the IL-1 receptor antagonist, anakinra, attenuates muscle wasting and adipose tissue browning in CKD mice via multiple cellular mechanisms (**Figure 8**). Administration of anakinra may represent a novel targeted treatment for cachexia in CKD patients, reversing muscle wasting and adipose tissue browning, and potentially improving long term outcomes in physical functioning, quality of life and survival.

**Figure 8:**
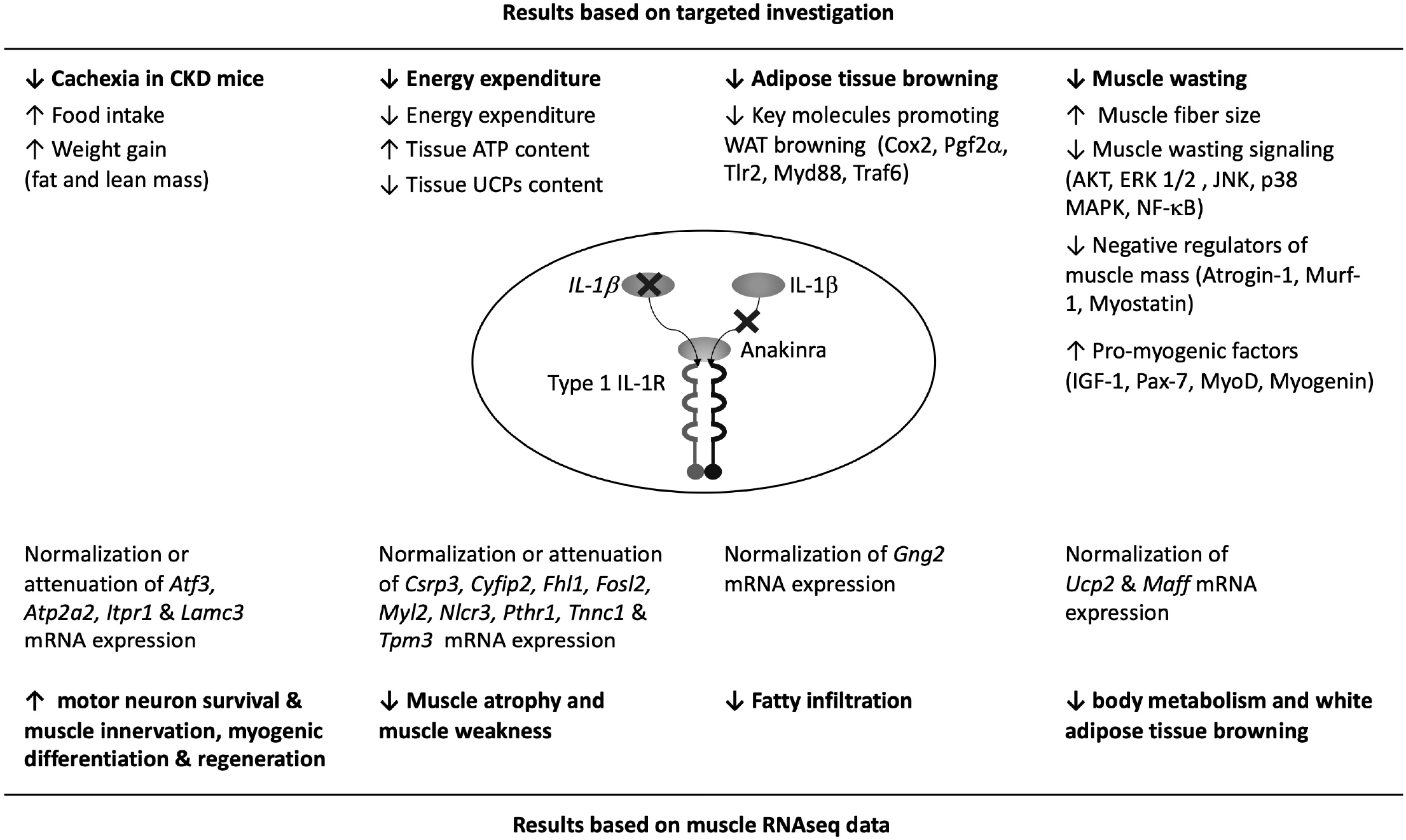
Summary of the beneficial effects of anakinra on cachexia, energy homeostasis, muscle wasting and adipose tissue browning in CKD mice.

## MATERIALS AND METHODS

### Study design

Wild-type (WT), *Il6*^−/−^, *Tnfα*^−/−^ and *Il1β*^−/−^ mice were on the same c57BL/6 genetic background. Six-week old male mice were used for the study. CKD was surgically induced by 2-stage 5/6 nephrectomy in WT, *Il6*^−/−^, *Tnfα*^−/−^ and *Il1β*^−/−^ mice while sham operation was performed in respective control mice. We have performed the following 5 studies. The study period was 6 weeks in all studies. **Study 1** – We measured gastrocnemius muscle IL-6, TNF-α and IL-1β mRNA and protein content in WT/CKD mice and pair-fed WT mice. Results were presented in Figure 1, A to F. **Study 2** – We evaluated the metabolic effects of genetic deletion of *Il6, Tnfα* and *Il1β* in CKD mice. Specifically, we compared *ad libitum* food intake and weight change in WT/CKD, *Il6*^−/−^/CKD, *Tnfα*^−/−^/CKD and *Il1β*^−/−^/ CKD mice relative to their respective sham controls. Results were shown in Figure 1, G & H. **Study 3** – We evaluated the beneficial effects of genetic deletion of *Il6, Tnfα* and *Il1β* in CKD mice beyond nutritional effects by employing a pair-feeding strategy. WT/CKD mice were fed *ad libitum* and then WT/sham mice as well as *Il6*^−/−^/CKD, *Tnfα*^−/−^/CKD and *Il1β*^−/−^/CKD mice and their respective sham controls were fed the same amount of rodent diet based on the recorded food intake of WT/CKD mice. Results were shown in Figure 1, I to N. **Study 4** - We evaluated the effects of anakinra in WT/CKD mice. WT/CKD and WT/Sham mice were given anakinra (2.5 mg/kg.day, IP) or vehicle (normal saline), respectively. All mice were fed *ad libitum*. We compared food intake and weight change in all groups of mice. Results were shown in Figure 2, A & B. **Study 5** - We evaluated the metabolic effects of anakinra in WT/CKD mice beyond nutritional stimulation by employing the pair-feeding strategy. WT/CKD and WT/Sham mice were given anakinra (2.5 mg/kg.day, IP) or vehicle (normal saline), respectively. Vehicle-treated WT/CKD mice were fed *libitum* while all other group of mice were fed the same amount of rodent diet based on the recorded food intake of vehicle-treated WT/CKD mice. Results were shown in Figure 2, E to J & Figures, 3 to 7.

### Body composition, metabolic rate and *in vivo* muscle function

Body composition was measured by quantitative magnetic resonance analysis (EchoMRI-100^™^, Echo Medical System). Twenty-four-hour metabolic rate (VO2) was measured using Oxymax indirect calorimetry (Columbus Instrument). *in vivo* muscle function (grip strength and rotarod activity) in mice was assessed using a grip strength meter (Model 47106, UGO Basile) and rotarod performance tool (model RRF/SP, Accuscan Instrument), respectively.^65^

### Serum and blood chemistry

BUN and serum concentration of bicarbonate was measured (**Supplemental Table 4S**). Serum creatinine were analyzed by LC-MS/MS method.^66^

### Protein assay for muscle and adipose tissue

Gastrocnemius muscle, inguinal white adipose tissue (WAT) and intercapsular brown adipose tissue (BAT) were processed in a homogenizer tube (USA Scientific, catalog 1420-9600) containing ceramic beads (Omni International, catalog 19-646) using a Bead Mill Homogenizer (Omni International). Protein concentration of tissue homogenate was assayed using Pierce BAC Protein Assay Kit (Thermo Scientific, catalog 23227). Uncoupling protein (UCP) protein content as well as adenosine triphosphate (ATP) concentration in adipose tissue and muscle homogenates were assayed. Protein contents of CD137, Tmem26, Tbx-1, Cox2, Pgf2a, Tlr2, Myd88, Traf6 in adipose tissue homogenates and protein contents of IL-6, TNF-α, IL-1β, phospho-Akt (pS473) and total Akt, phospho-ERK 1/2 (Thr202/Ty2r204) and total ERK 1/2, phospho-JNK (Thr183/Tyr185) and total JNK, phospho-p38 MAPK (Thr180/Tyr182) and total p38 MAPK, NF-κB p50 (phospho-Ser337) and total NF-κB p50, NF-κB p65 (phospho-Ser536) and total NF-κB p65, Iκκa (phosphor-Ser536) and total Iκκa in muscle homogenates were assayed (**Supplemental Table 4S**).

### Gastrocnemius weight, fiber size and fatty infiltration

Left gastrocnemius was excised and weighted. Fibre cross-sectional areas of left gastrocnemius were measured using ImageJ software (https://rsbweb.nih.gob/ij/).^12^ In addition, portion of dissected right gastrocnemius muscle samples were incubated with Oil Red O (Oil Red O Solution, catalog number O1391-250 ml, Sigma Aldrich). Detailed procedures for Oil Red O staining were in accordance with published protocol.^67^ We followed a recently established protocol to quantify muscle fat infiltration. Acquisition and quantification of images were analyzed using ImageJ software.^68^

### Muscle RNAseq analysis

We performed RNAseq analysis on gastrocnemius muscle mRNA in 12-month old WT/CKD mice versus age-appropriate WT/Sham mice. Detailed procedures for mRNA extraction, purification and subsequent construction of cDNA libraries as well as analysis of gene expression was published.^12^ We then performed Ingenuity Pathway Analysis enrichment tests for 20 previously identified differentially expressed muscle genes in WT/CKD mice versus WT/Sham mice, focusing on pathways related to energy metabolism, skeletal and muscle system development and function, and organismal injury and abnormalities. In this study, we performed qPCR analysis for these top 20 differentially expressed gastrocnemius muscle genes in pair-fed WT/CKD and WT/Sham mice treated with anakinra or vehicle, respectively.

### Quantitative real-time PCR

Total RNA from adipose and gastrocnemius muscle samples were isolated using TriZol (Life Technology) and reverse-transcribed with SuperScript III Reverse Transcriptase (Invitrogen). Quantitative real-time RT-PCR of target genes were performed using KAPA SYBR FAST qPCR kit (KAPA Biosystems). Expression levels were calculated according to the relative 2^-ΔΔCt^ method.^65^ All primers for target genes are listed (**Supplemental Table 5S**).

### Statistics

Continuous variables are expressed as mean ± S.E.M. We assessed the statistical significance of differences between groups using two-sample t-tests. All tests were two-sided. A p value less than 0.05 was considered significant. Statistical analyses were performed using SPSS software version 16.0 for Macintosh.

## Ethical statement

This study was performed in strict accordance with the recommendations in the Guide for the Care and Use of Laboratory Animals of the National Institutes of Health. All mice were handled according to approved institutional animal care and use committee (IACUC) protocols (S07154) of the University of California, San Diego.

## Author contribution statement

WWC, HMH and RHM conceived the study and designed the experiments. WWC, RZ, SH, ZW, AZ and PZ performed experiments. WWC and RHM analyzed the data. WWC, HMH and RHM interpreted the results and wrote the manuscript. All authors reviewed and approved the manuscript.

## Acknowledgements

We thank Dr. Jianhua Shao, UCSD Pediatric Diabetes Research Center for the use of EchoMRI-100^™^. The Moores Cancer Center Tissue Technology Shared Resource at University of California San Diego is supported by a National Cancer Institute Cancer Center Support Grant (CCSG Grant P30CA23100). The Acute Kidney Injury Research Bioanalytical Core at the O’Brien Center of the University of Alabama at Birmingham is supported by a P30 grant (DK 079337) from the National Institute of Diabetes and Digestive and Kidney Diseases (NIDDK).

---

**Supplemental Table 1S:** Serum and blood chemistry of CKD and control mice. CKD in wild type (c57BL/6J) mice was induced by 2-stage 5/6 nephrectomy and sham operation was performed in control mice. Mice were fed *ad libitum*. Mice were sacrificed at the end of 6 weeks study and serum and blood chemistry was measured. Data are expressed as mean ± SEM. Result of serum chemistry of WT/CKD mice were compared to WT/sham mice. # p<0.05, significantly increased in WT/CKD mice relative to WT/sham mice.

|  | WT/sham<br>n = 12 |  |  | WT/CKD<br>n = 12 |  |  |
| --- | --- | --- | --- | --- | --- | --- |
| BUN (mg/dL) | 35.4 | ± | 6.7 | 73.5 | ± | 12.5 # |
| Creatinine (mg/dL) | 0.08 | ± | 0.02 | 0.24 | ± | 0.02 # |
| Bicarbonate (mmol/L) | 26.5 | ± | 2.3 | 26.8 | ± | 3.7 |
| PTH (pg/mL) | 123.6 | ± | 21.5 | 278.6 | ± | 23.2 # |

**Supplemental Table 2S:**
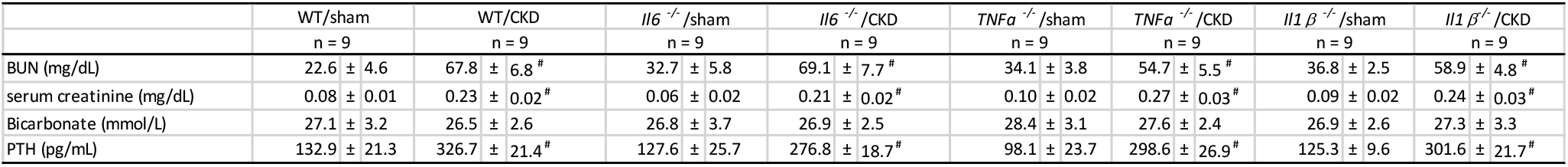
Serum and blood chemistry of *Il6*^−/−^, *Tnfα*^−/−^, *Il1β*^−/−^ and wild type control mice. Genetic background of *Il6*^−/−^, *Tnfα*^−/−^, *Il1β*^−/−^ and wild type control mice were c57BL/6J. CKD in *Il6*^−/−^, *Tnfα*^−/−^, *Il1β*^−/−^ and WT mice were surgically induced by 5/6 nephrectomy while sham operation was performed in respective control mice. Mice were fed *ad libitum* and *e*xperiment period was 6 weeks. Data are expressed as mean ± SEM. Result of *Il6*^−/−^, *Tnfα*^−/−^, *Il1β*^−/−^ mice were compared to WT mice. # p<0.05, significantly increased in *Il6*^−/−^, *Tnfα*^−/−^ and *Il1β*^−/−^ mice relative to WT/sham mice.

|  | WT/sham |  | WT/CKD |  | *Il6*<sup>-/-</sup>/sham |  | *Il6*<sup>-/-</sup>/CKD |  | *Tnfα*<sup>-/-</sup>/sham |  | *Tnfα*<sup>-/-</sup>/CKD |  | *Il1β*<sup>-/-</sup>/sham |  | *Il1β*<sup>-/-</sup>/CKD |  |
| --- | --- | --- | --- | --- | --- | --- | --- | --- | --- | --- | --- | --- | --- | --- | --- | --- |
|  | n = 9 |  | n = 9 |  | n = 9 |  | n = 9 |  | n = 9 |  | n = 9 |  | n = 9 |  | n = 9 |  |
| BUN (mg/dL) | 22.6 | ± 4.6 | 67.8 | ± 6.8<sup>#</sup> | 32.7 | ± 5.8 | 69.1 | ± 7.7<sup>#</sup> | 34.1 | ± 3.8 | 54.7 | ± 5.5<sup>#</sup> | 36.8 | ± 2.5 | 58.9 | ± 4.8<sup>#</sup> |
| serum creatinine (mg/dL) | 0.08 | ± 0.01 | 0.23 | ± 0.02<sup>#</sup> | 0.06 | ± 0.02 | 0.21 | ± 0.02<sup>#</sup> | 0.10 | ± 0.02 | 0.27 | ± 0.03<sup>#</sup> | 0.09 | ± 0.02 | 0.24 | ± 0.03<sup>#</sup> |
| Bicarbonate (mmol/L) | 27.1 | ± 3.2 | 26.5 | ± 2.6 | 26.8 | ± 3.7 | 26.9 | ± 2.5 | 28.4 | ± 3.1 | 27.6 | ± 2.4 | 26.9 | ± 2.6 | 27.3 | ± 3.3 |
| PTH (pg/mL) | 132.9 | ± 21.3 | 326.7 | ± 21.4<sup>#</sup> | 127.6 | ± 25.7 | 276.8 | ± 18.7<sup>#</sup> | 98.1 | ± 23.7 | 298.6 | ± 26.9<sup>#</sup> | 125.3 | ± 9.6 | 301.6 | ± 21.7<sup>#</sup> |

**Supplemental Table 3S:** Serum and blood chemistry of CKD and wild type control mice. WT/CKD and WT/sham mice were given anakinra (2.5 mg/kg.day, IP) or vehicle (normal saline), respectively. Vehicle-treated WT/CKD mice were fed *ad libitum* while all other group of mice were fed the same amount of rodent diet based on the recorded food intake of vehicle-treated CKD mice. The study period was 6 weeks. Result of serum chemistry of WT/CKD+Vehicle mice were compared to WT/sham+Vehicle mice while results of WT/CKD+Anakinra mice were compared to WT/sham+Anakinra. Data are expressed as mean ± SEM. # p<0.05, significantly increased in WT/CKD+Vehicle and WT/CKD+Anakinra mice relative to WT/sham+Vehicle and WT+Anakinra mice, respectively.

|  | WT/sham+Vehicle |  |  | WT/sham+Anakinra |  |  | WT/CKD+Vehicle |  |  | WT/CKD+Anakinra |  |  |
| --- | --- | --- | --- | --- | --- | --- | --- | --- | --- | --- | --- | --- |
|  | n = 9 |  |  | n = 9 |  |  | n = 9 |  |  | n = 9 |  |  |
| BUN (mg/dL) | 26.7 | ± | 4.7 | 36.1 | ± | 3.8 | 78.3 | ± | 9.3 # | 65.8 | ± | 3.6 # |
| serum creatinine (mg/dL) | 0.06 | ± | 0.01 | 0.09 | ± | 0.02 | 0.25 | ± | 0.02 # | 0.19 | ± | 0.03 # |
| Bicarbonate (mmol/L) | 27.6 | ± | 2.5 | 27.1 | ± | 3.1 | 26.7 | ± | 2.5 | 27.5 | ± | 4.4 |
| PTH (pg/mL) | 108.7 | ± | 10.6 | 109.6 | ± | 11.6 | 287.6 | ± | 7.9 # | 298.4 | ± | 11.6 # |

**Supplemental Table 4S:** Immunoassay information for blood and serum chemistry, muscle adenosine triphosphate content as well as muscle and adipose tissue protein analysis.

| <u>Blood & Serum chemistry</u> | <u>Assay information</u> |
| --- | --- |
| Bicarbonate & BUN | VetScan Comprehensive Diagnostic Profile, Abaxis, 500-0038 |
| Creatinine | LC-MS/MS method |
| Mouse PTH 1-84 ELISA Kit | Immutopics, 60-2305 |
| <u>Muscle & adipose tissue</u> | <u>Assay information</u> |
| ATP Assay Kit (Colorimetric / Fluorometric) | Abcam, ab83355 |
| Mouse CD137 (TNFRSF9) ELISA kit | LifeSpan BioSciences, LS-F2852-1 |
| Mouse COX2 ELISA kit | LifeSpan BioSciences, LS-F37124-1 |
| Mouse IL-1 beta ELISA Kit | RayBiotech, ELM-IL1b-CL |
| Mouse IL-6 ELISA Kit | RayBiotech, ELM-IL6 |
| Mouse TNF-alpha ELISA Kit | RayBiotech, ELM-TNFalpha |
| Human/mouse/rat Phospho-AKT (S473) and total AKT ELISA | Raybiotech, PEL-AKT-S473-T-1 |
| Mouse, rat, human ERK1/2, JNK, p38 MAPK phosphorylation ELISA sampler kit | RayBiotech, CBEL-ERK-SK |
| Mouse IKK-alpha (phospho-Thr23) ELISA kit | LifeSpan BioSciences, LS-F1537-1 |
| Mouse MYD88 ELISA kit | LifeSpan BioSciences, LS-F33219-1 |
| Mouse NFkB p50 (phospho-Ser337) ELISA kit | Aviva Systems Biology, OKAG00322 |
| Mouse NFkB p65 (phospho-Ser536) ELISA kit | RayBiotech, PEL-NFKBP65-S536-T-1 |
| Mouse total NFkB p65 ELISA kit | RayBiotech, PEL-NFKBP65-S536-T-1 |
| Mouse Prostaglandin F2 alpha ELISA kit | MyBioSource, MBS266867 |
| Mouse TBX1 ELISA kit | Antibodies-online, ABIN6221756 |
| Mouse TLR2 ELISA kit | Abcam, ab224880 |
| Mouse TNF-alpha ELISA Kit | RayBiotech, ELM-TNFalpha |
| Mouse TMEM26 ELISA kit | MyBiosource, MBS9319338 |
| Mouse TRAF6 ELISA kit | LifeSpan BioSciences, LS-F53451 |
| Mouse Ucp1 ELISA kits | Aviva Systems Biology, OKCD02970 |
| Mouse Ucp3 ELISA kits | Aviva Systems Biology, OKEH05259 |

**Supplemental Table 5S:**
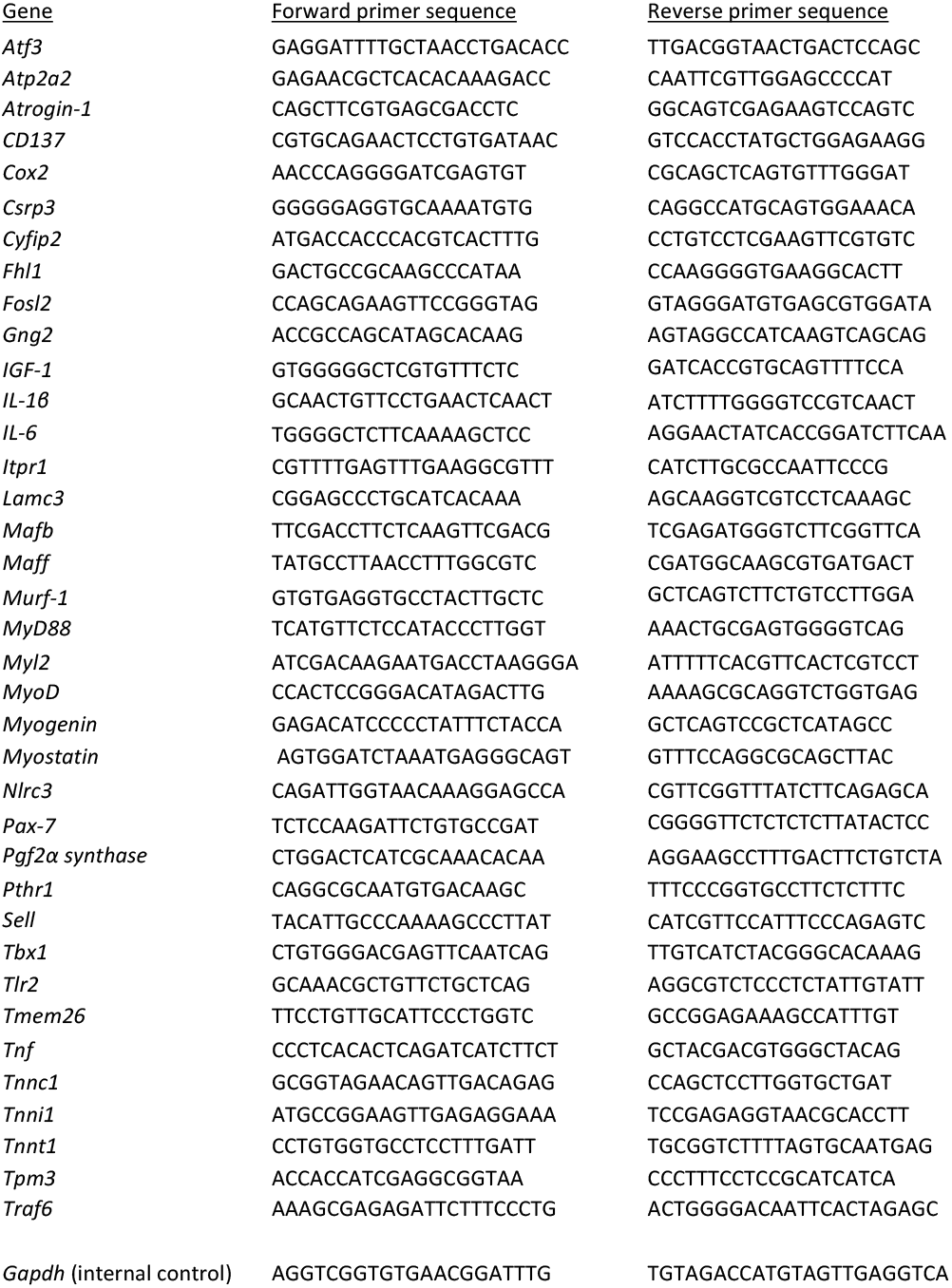
PCR primer information.

**Supplemental Figure 1:**
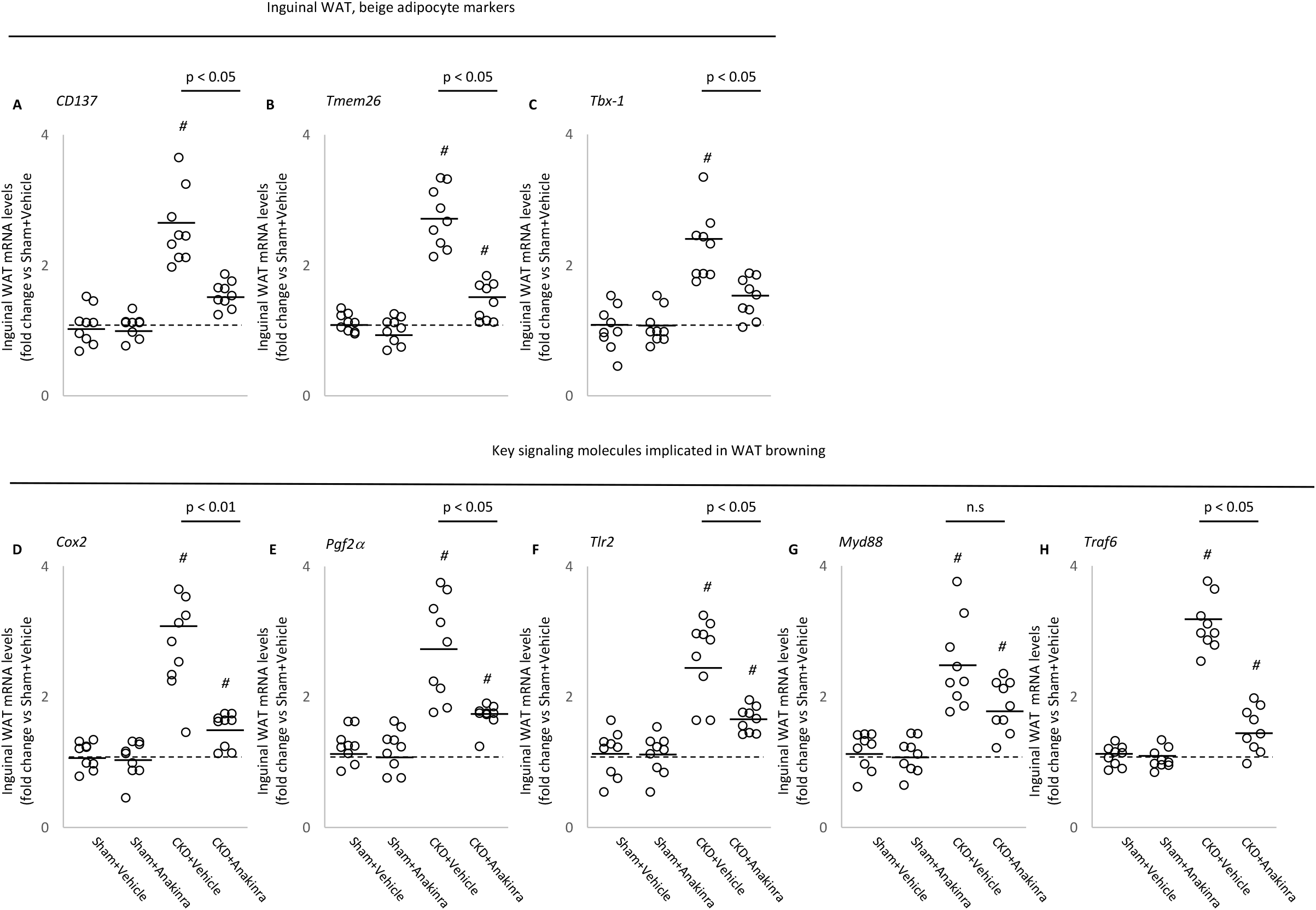
Anakinra attenuates adipose tissue browning in CKD mice. Gene expression in inguinal white adipose tissue. Gene expression of beige adipocyte markers (*CD137, Tmem26* and *Tbx-1*) in inguinal white adipose tissue was measured by qPCR. In addition, gene expression of Cox2 signaling pathway (*Cox2* and *Pgf2a*) and toll like receptor pathway (*Tlr2, MyD88* and *Traf6*) in inguinal white adipose tissue was measured by qPCR. Final results were expressed in arbitrary units, with one unit being the mean level in WT/sham+Vehicle mice. Data are expressed as mean±SEM. Results of WT/CKD+Vehicle were compared to WT/sham+Vehicle and WT/CKD+Anakinra were compared to WT/sham+Anakinra, respectively. In addition, results of WT/CKD+Anakinra were also compared to WT/CKD+Vehicle. # p < 0.05.

**Supplemental Figure 2:**
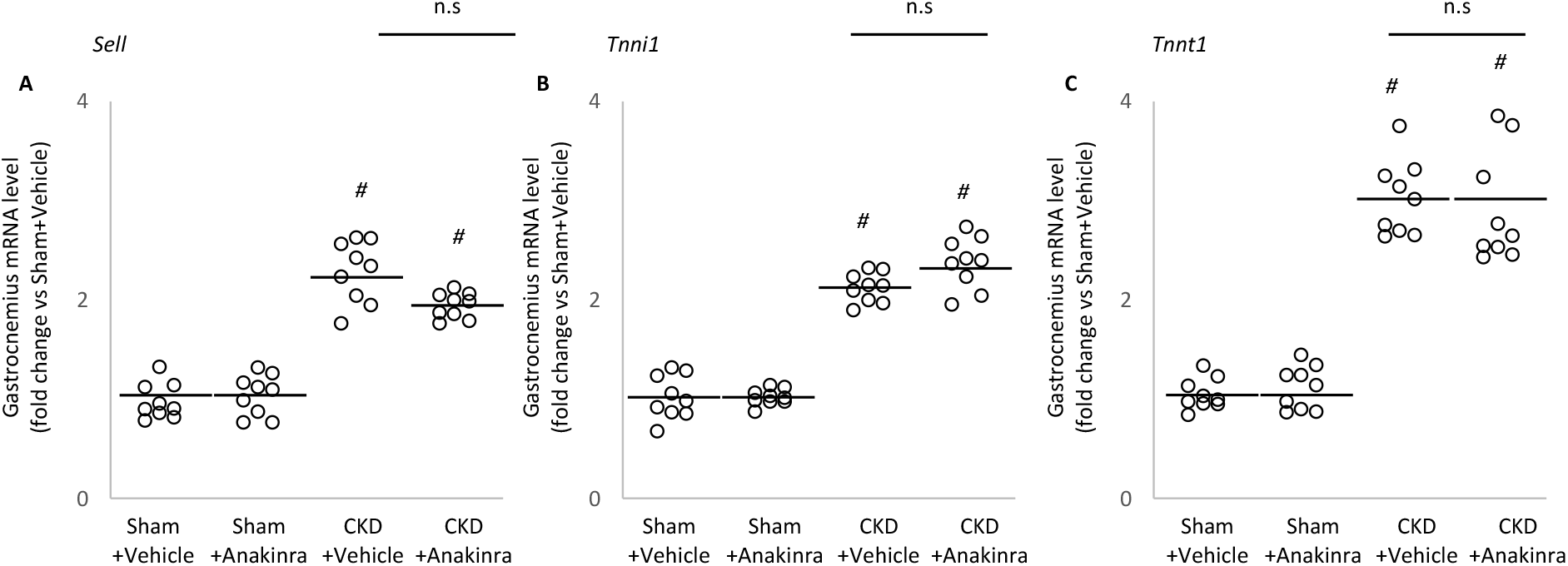
Differential expression muscle genes in CKD mice versus control mice. Gastrocnemius muscle expression of interested genes (*Sell, Tnni1* and *Tnnt1*) in mice was measured by qPCR. Final results were expressed in arbitrary units, with one unit being the mean level in vehicle-treated WT/Sham mice. Results are analyzed and expressed as in Supplemental Figure 1.

